# Extracellular injection system combined with peptides for intracellular *Staphylococcus aureus* treatment

**DOI:** 10.64898/2026.07.06.736670

**Authors:** Lei Feng, Yingying Qiao, Haishan Xu, Gaojie Wang, Shilong Ren, Xudong Ouyang, Ningning Song, Xiangxiang Zhao, Xianchao Feng

## Abstract

The inaccessibility of intracellular bacteria has long rendered the treatment of Staphylococcus aureus infections an challenge. Studies have demonstrated that the extracellular injection system PVC can accurately deliver proteins into cells, which would not need small molecules, and enables effective intracellular delivery of antimicrobial peptides for treatment. Accordingly, we selected antimicrobial peptides including Cecropin, LL37 and Indolicidin that possess potent bactericidal activity, and established the Directed Antimicrobial Assault platform (DAAT) by leveraging the intracellular delivery capacity of PVC. DAAT Cecropin, DAAT LL37 and DAAT Indolicidin inhibited intracellular bacteria in a dose-dependent manner, with DAAT LL37 reaching 86.76% inhibition; after 72 h of treatment, viable-cell numbers reduse to 66—82-fold those of the control. Tail-fibre retargeting enabled direct extracellular *S. aureus* killing, while combined DAAT therapy promoted wound healing in mice. These findings expand the utility of PVC-derived nanosyringes and establish DAAT as a modular platform for intracellular antimicrobial peptide therapy.

## 1. Introduction

The central challenge in treating clinical intracellular bacterial infections is the difficulty of delivering antibacterial agents effectively into cells, particularly in the context of intracellular invasion by methicillin-resistant *Staphylococcus aureus* (MRSA) (Peyrusson et al., 2020; Hommes et al., 2022). *S. aureus* can actively invade host cells and persist by exploiting the intracellular environment as a protective barrier, ultimately causing persistent infection (Peyrusson et al., 2020; Volk et al., 2024). Although next-generation antibiotics, antimicrobial nanomaterials, phage-derived enzymes and anti-virulence strategies exhibit bactericidal activity against extracellular MRSA, their activity against intracellular bacteria remains limited (de Oliveira et al., 2024). Antimicrobial peptides (AMPs) are highly promising anti-*S. aureus* candidates owing to their rapid, multi-target mechanisms of action; however, their susceptibility to proteolysis, inefficient transmembrane penetration and subtherapeutic accumulation at infection sites restrict their intracellular application (Brogden, 2005; Cesaro et al., 2023; de Oliveira et al., 2024). Therefore, a safe and efficient delivery system is urgently needed to deliver AMPs directly to intracellular infection sites.

Extracellular contractile injection systems, particularly the bacterium-derived *Photorhabdus* virulence cassette (PVC), can deliver protein cargos into target cells through a self-powered contraction mechanism, and its confined tubular compartment provides a protective space that is well suited for antimicrobial peptide (AMP) loading and delivery (Vlisidou et al., 2019; Desfosses et al., 2019; Jiang et al., 2022; Kreitz et al., 2023). Conventional tags ease AMP expression at the cost of activity, as they can obscure cationic and amphipathic motifs and impose additional cleavage–repurification steps (Li, 2009; Li, 2011; Sadeeq et al., 2025). In this context, the Pdp1-guided PVC loading strategy offers a integrated solution: Pdp1 transiently masks the exposed cationic features of AMPs during cytosolic expression, limits premature toxicity to the production host, and then directs the peptide cargo into the PVC inner tube during particle assembly (Jiang et al., 2022). Once enclosed within the PVC lumen, AMPs are physically shielded until contractile injection into target cells, thereby coupling host-tolerant expression, protective encapsulation and intracellular delivery in one biosynthetic platform for *S. aureus* therapy (Vlisidou et al., 2019; Jiang et al., 2022; Kreitz et al., 2023).

Treatment of *Staphylococcus aureus* infection cannot rely solely on eliminating intracellular bacterial reservoirs; the large extracellular bacterial population and its potential to drive reinvasion and infection recurrence must also be considered (Peyrusson et al., 2020; Hommes et al., 2022; Volk et al., 2024). Coordinated clearance of both intracellular and extracellular *S. aureus* therefore represents the optimal strategy. However, native PVC lacks an intrinsic recognition module for directly targeting *S. aureus*. Previous studies have shown that PVC tropism can be reprogrammed through tail-fibre modification, providing a foundation for overcoming this limitation (Kreitz et al., 2023). The receptor-binding protein Gp45 from *S. aureus* phage φ11 provides a rational targeting element: it mediates host recognition through GlcNAc-modified wall teichoic acid and functions as a key baseplate-associated receptor-binding protein (Xia et al., 2011; Li et al., 2016; Koç et al., 2016). Therefore, introducing a Gp45-derived recognition domain into the PVC tail fibre can confer the ability to bind directly to *S. aureus*, anchor to its surface and execute contractile injection. This modification not only enables direct killing of extracellular *S. aureus*, but also overcomes the previous inability of PVC to target bacterial cells, thereby extending DAAT into a coordinated intracellular and extracellular antibacterial platform.

In this study, we developed Directed Antimicrobial Assault Technology (DAAT), a PVC-derived contractile nanosyringe platform that integrates intracellular AMP delivery with extracellular *S. aureus* targeting. For intracellular therapy, Cecropin, LL-37 or Indolicidin was fused to a Pdp1-derived loading module and recruited into the PVC inner tube during particle assembly, enabling host-tolerant AMP expression, protective encapsulation and contractile injection. Successful AMP loading was validated by nanosyringe morphology, PVC structural protein profiling, HiBiT luminescence and low-molecular-weight gel analysis. DAAT suppressed intracellular *S. aureus* in a dose-dependent manner, with DAAT-LL37 showing the strongest activity; 256 ng μL⁻¹ preserved HT-29 compatibility, whereas 512 ng μL⁻¹ induced host-cell stress. For extracellular clearance, we further propose retargeting the PVC tail fibre with the *S. aureus* phage φ11 receptor-binding protein Gp45, enabling recognition of GlcNAc-modified wall teichoic acid, bacterial anchoring and contractile injection. Ultimately, intracellular–extracellular synergistic antibacterial activity mediated by PVC technology promoted the healing of infected skin tissues in mice. Thus, DAAT provides a dual-mode strategy against intracellular reservoirs and extracellular *S. aureus*.

## 2. Materials and Methods

### 2.1 Evaluation of Antibacterial Activity of Synthetic Antimicrobial Peptides

Cecropin, LL-37, and Indolicidin were synthesized by Tsingke Biotechnology Co., Ltd. (Beijing, China). To further evaluate the in vitro antibacterial activity of the three antimicrobial peptides, a concentration-gradient plate-counting assay was performed. Each peptide was prepared at serial concentrations of 0, 1, 2, 4, 8, 16, 32, 64, 128, 256, and 512 ng μL⁻¹ and incubated with the tested bacterial suspension. After incubation, the treated bacterial suspensions were serially diluted and spread onto solid agar plates. Colony formation was observed after cultivation to assess the inhibitory effects of different peptide concentrations on bacterial survival and proliferation.

### 2.2 Reagents and plasmid construction

The PVCpnf structural plasmid containing *pvc1–16*, the payload plasmid encoding *Pdp1/Pnf*, and the regulatory plasmid carrying PAU_RS16570–RS24015 were purchased from Addgene (198291, 198287, and 198317). Antimicrobial peptide coding sequences and all oligonucleotides required for cloning were synthesized by General Biotech Co., Ltd. (China). For plasmid construction, the pPayload backbone and corresponding insert fragments were amplified by PCR using Phusion Flash 2× Master Mix (Thermo Fisher Scientific, F548S), with the extension step set at 15 s. The plasmid construction involve uses T5 exonuclease-dependent assembly master mix (TEDA; Laboratory-prepared referenced to a previously reported; 30 ℃ for 40 min) or 2× Ezmax® Ultra Universal CloneMix (TOLOBIO, China; 50℃ for 30 min). The assembled products were then transformed into chemically competent *E. coli* DH5α cells.

For tail-fibre retargeting, the PVCpnf structural plasmid containing *pvc1–16*. The original AD5RGDPK7 targeting sequence within Pvc13 was replaced with the Gp45 coding sequence to confer recognition of *Staphylococcus aureus*. The Pvc13-containing plasmid backbone and the Gp45 insert were amplified with primers carrying the required homologous overlaps and assembled by homologous recombination. Plasmid reconstruction was performed using the same assembly systems described above. The assembled products were introduced into electrocompetent *E. coli* EPI300 cells by electroporation, and positive clones were identified by colony PCR and confirmed by DNA sequencing.

### 2.3 DAAT expression and purification

DAAT expression and purification were performed with minor modifications to the protocol described by Kreitz et al (2023). Briefly, the constructed antimicrobial peptide–encoding pPayload plasmid and one corresponding variant of the PVC structural plasmid (pPVC) were electroporated into the same electrocompetent Escherichia coli EPI300 cells. Positive clones were cultured in LB medium (Beyotime, ST163) supplemented with kanamycin (Beyotime, ST101; 50 μg/mL, 3 μL) and ampicillin (Beyotime, ST008; 100 μg/mL, 3 μL) at 37°C and 200 rpm for 12 h. The overnight culture was then diluted 1:100 into 500 mL of fresh LB medium containing the same antibiotics and further incubated at 30 °C and 160 rpm for 24 h. Cells were harvested by centrifugation at 4,000 × g for 10 min at 4 °C. The resulting cell pellets were either processed immediately or stored at −80 °C for 1–2 months before use. For DAAT purification, the collected cell pellets were first resuspended in lysis buffer prepared exactly as described by Kreitz et al (2023). A protease inhibitor cocktail (Roche, 11873580001; 1×), DNase I (Thermo Fisher Scientific, EN0521; 25 μg/mL), and lysozyme (Sigma-Aldrich, L6876; 100 μg/mL) were then added, followed by gentle rotation at 25°C for 1.5 h. The lysate was clarified by centrifugation at 4,000 × g for 30 min at 4°C, and the resulting supernatant was subjected to ultracentrifugation at 120,000 × g for 2 h at 4°C. The pellet was resuspended in PBS (Life Technologies, 10010049) and further clarified by centrifugation at 16,000 × g for 15 min at 4°C. To improve sample purity, the ultracentrifugation–resuspension cycle was repeated twice. Finally, the clarified suspension was ultracentrifuged once more at 120,000 × g for 2 h at 4°C to obtain highly purified DAAT particles.

The final DAAT pellet was resuspended in 200 μL of phosphate buffer at pH 7.5, and the DAAT concentration was determined from the A280 value using an Implen NanoPhotometer N80 spectrophotometer (Implen, Germany). Purified DAAT preparations were stored at 4°C for 7 days, or in phosphate buffer containing 10% glycerol at −20°C for 3–4 months.

### 2.4 Bacterial strain and culture

The GFP-expressing *Staphylococcus aureus* strain USA300/Eno-GFP was kindly provided by the research group of Prof. Xiancai Rao at Army Medical University. In this strain, the GFP-encoding sequence was integrated into the bacterial genome, allowing bacterial cells to be visualized as green fluorescent signals under fluorescence microscopy. For routine propagation, USA300/Eno-GFP was cultured in tryptic soy broth (TSB) at 37 °C with shaking at 200 rpm overnight. Before use in infection-related experiments, the overnight culture was refreshed in sterile TSB medium and adjusted to the required bacterial density according to the experimental design.

### 2.5 Cell culture

HT-29 cells were purchased from Procell Life Science & Technology Co., Ltd (China). Cell culture and routine passaging were performed according to the supplier’s instructions. The cell lines used in this study were purchased from commercial suppliers and were not further authenticated or tested for mycoplasma contamination before experimental use. Briefly, cells were maintained in T25 culture flasks (Corning, 430639) at 37 °C in a humidified incubator containing 5% CO₂ (Heracell™ 150i; Thermo Scientific, USA). Complete culture medium supplied by Procell was used for routine maintenance. For infection-model establishment and related assays requiring antibiotic-free conditions, cells were cultured in DMEM/F-12 (1:1) medium (Purino, PM150312) supplemented with 10% fetal bovine serum (Martínez-Maqueda et al., 2015).

### 2.6 Establishment of an intracellular *Staphylococcus aureus* infection model

To establish an intracellular *Staphylococcus aureus* infection model, HT-29 cells were seeded into 6-well plates at a density of 1 × 10⁵–1 × 10⁶ cells per well in 2 mL of complete culture medium and incubated for 24 h at 37 °C in a humidified atmosphere containing 5% CO₂ to allow cell attachment and recovery. *S. aureus* suspensions were then added to each well at 1.6 × 10⁴, 3.2 × 10⁴, 6.4 × 10⁴, and 1.28 × 10⁵ CFU per well, respectively, followed by incubation for 1 h to permit bacterial adhesion and internalization (Hess et al., 2003). HT-29 cell numbers were determined using a hemocytometer, whereas bacterial concentrations were estimated by *OD₆₀₀* measurement and converted to CFU values based on a pre-established *OD₆₀₀*–CFU standard curve (Stevenson et al., 2016). Bacterial concentrations were estimated by *OD₆₀₀*measurement and converted to CFU values using a pre-established *OD₆₀₀*–CFU standard curve. The bacterial inoculum was calculated as CFU per well = CFU mL⁻¹ × V, where V represents the volume of bacterial suspension added to each well in millilitres. After infection, the culture medium was removed, and the cells were gently washed twice with sterile PBS to eliminate non-adherent bacteria. Subsequently, 2 mL of fresh complete medium containing gentamicin at 50 ng/μL and penicillin at 10 ng/μL was added to each well to kill residual extracellular *S. aureus*, and the cells were further incubated for 2 h under standard culture conditions (Rollin et al., 2017; Kim et al., 2019). The antibiotic-containing medium was then discarded, and the cells were washed twice again with sterile PBS. Fresh complete medium supplemented with gentamicin at 10 ng/μL was added to suppress extracellular bacterial regrowth during the subsequent incubation period. After an additional 6 h of culture, cell morphology, adherence, and overall growth status were examined microscopically. Based on the balance between effective intracellular infection and preservation of host-cell viability, 6.4 × 10⁴ CFU per well was selected as the optimal bacterial inoculum for subsequent experiments.

### 2.7 Electron microscopy

Negative-stain transmission electron microscopy (TEM) of purified DAAT particles was performed with technical support from Scientific Compass (Beijing, China), an independent testing laboratory accredited under ISO/IEC 17025. Briefly, 10 μ L of each sample was adsorbed onto plasma-cleaned 200-mesh carbon-coated copper grids. The grids were then negatively stained with 2% uranyl acetate using an Electron Microscopy Sciences C1430 system. Imaging was conducted on a JEOL JEM-2100F transmission electron microscope equipped with a Gatan US1000 CCD camera under low-dose conditions at an accelerating voltage of 120 kV. Representative fields were recorded for subsequent morphological analysis (Booth et al., 2011).

### 2.8 SDS-PAGE analysis

The protein profile of purified DAAT samples was examined by sodium dodecyl sulfate–polyacrylamide gel electrophoresis (SDS-PAGE) (Lim et al., 2023; Kilic et al., 2024). Briefly, 20 μL of each DAAT preparation was mixed with 5× SDS-PAGE loading buffer at a 4:1 volume ratio. The mixture was gently pipetted to ensure complete homogenization and then heated at 98℃ for 10 min in a dry bath to denature the proteins before loading. Electrophoresis was performed using a discontinuous polyacrylamide gel system consisting of a 5% stacking gel and a 12% resolving gel. After the gel was assembled in the electrophoresis tank, 1× SDS-PAGE running buffer was added. A pre-stained protein molecular weight marker was loaded at 10 μL per lane, and each denatured DAAT sample was loaded at 20 μL per lane. Electrophoresis was initially carried out at 80 V until the samples migrated through the stacking gel, after which the voltage was increased to 120 V and maintained until the bromophenol blue dye front approached the bottom of the gel. Following electrophoresis, the gel was immersed in Coomassie Brilliant Blue staining solution for 30 min (Lim et al., 2023). The stained gel was subsequently destained until the background became clear and distinct protein bands were visible. Gel images were acquired using a gel documentation system. The major protein bands in the DAAT preparations were evaluated by comparison with the molecular weight marker to assess the protein composition and band distribution of the purified particles.

### 2.9 Verification of antimicrobial peptide loading

To further confirm the successful loading of antimicrobial peptides into DAAT particles, a HiBiT tag was individually fused to the C-terminus of each antimicrobial peptide. The signal intensity was then measured using the Nano-Glo® HiBiT Extracellular Detection System (Promega, NG2421) (Promega, 2017). HiBiT is a bioluminescence-based protein detection strategy that enables highly sensitive quantification through the high-affinity complementation between the short HiBiT peptide tag and the LgBiT subunit (Schwinn et al., 2018; Boursier et al., 2020). All experimental procedures and data calculations were performed according to the manufacturer’s instructions. Each sample was analyzed in three independent technical replicates to minimize experimental variation and improve the reliability of the resulting conclusions.

### 2.10 Optical microscopy imaging

The establishment of the *Staphylococcus aureus* infection model was evaluated by microscopic observation and colony-based bacterial enumeration. Normal HT-29 cells and infected HT-29 cells were examined by optical microscopy to compare cellular morphology, including, spreading, and infection-associated morphological changes. In parallel, the GFP fluorescence channel was used to visualize USA300/Eno-GFP bacteria and to assess the infection status of HT-29 cells. Representative bright-field and fluorescence images were acquired using 4× and 40× objectives, with identical imaging settings applied to comparable groups. To further determine the bacterial distribution under different infection inocula, HT-29 cells were infected with USA300/Eno-GFP at bacterial inputs of 1.6, 3.2, 6.4, 12.8, and 25.6 10⁴ CFU per well. After infection, extracellular and intracellular bacterial burdens were separately quantified by colony enumeration using a serial-dilution spot-plating assay (Thomas et al., 2015). For extracellular bacterial analysis, culture supernatants or extracellular fractions were collected after infection and diluted in sterile PBS or TSB medium. For intracellular bacterial recovery, infected cells were washed with sterile PBS to remove non-adherent bacteria, treated with antibiotic-containing medium to eliminate residual extracellular bacteria, and subsequently lysed to release internalized bacteria (Sharma & Puhar, 2019). The collected bacterial suspensions were serially diluted, spotted onto TSB agar plates, and incubated at 37 ° C until visible colonies formed. Colony numbers were recorded from countable dilution spots and used to compare extracellular and intracellular bacterial loads across the different infection doses. Pseudocolor optical microscopy was performed using an AJ-VERT optical microscopy imaging system equipped with AJ-VERT image acquisition software (AJ-VERT; AOR Industrial Co., Ltd., Shenzhen, China). Briefly, untreated control cells cultured overnight in 6-well plates, HT-29 cells after establishment of the intracellular *Staphylococcus aureus* infection model, and cells collected after one round of treatment were prepared directly in the culture plates for imaging. Before image acquisition, the culture medium was carefully aspirated, and the cells were gently rinsed twice with sterile PBS to remove residual medium and loosely attached materials. The plates were then immediately placed on the microscope stage and examined using a 20× objective lens. Image acquisition and pseudocolor adjustment were both carried out in the AJ-VERT software. The imaging parameters were kept as follows: display unit, μm; digital zoom, 67%; RGB channel values, red = 125, green = 164, and blue = 167; hue, +6; saturation, 255; brightness, 35; contrast, 100; gamma, 1.24; and flicker-suppression mode, direct current (DC). Clear fields containing normal and infected cells were first located through the eyepiece to allow reliable morphological discrimination between healthy and abnormal cells. Representative fields from all experimental groups were recorded using the same acquisition settings and pseudocolor calibration parameters to maintain consistency and allow direct comparison among samples.

### 2.11 Drug concentration-gradient screening

Drug concentration-gradient screening was performed by comparing the inhibitory effects of different DAAT treatments on the experimental groups and controls. HT-29 cells were seeded into 96-well cell culture plates with an appropriate volume of complete medium per well and allowed to attach uniformly. After stable cell adherence, the original medium was removed and replaced with antibiotic-free medium containing serial concentrations of DAAT-Cecropin, DAAT-LL37, or DAAT-Indolicidin. The final concentrations of each DAAT preparation were set at 512, 256, 128, 64, 32, 16, 8, 4, 2, and 1 ng μL⁻¹. After 24 h of treatment, the contents of each well were thoroughly mixed, and the optical density at 600 nm (*OD₆₀₀*) was measured using a microplate reader to estimate bacterial growth based on medium turbidity (Balouiri et al., 2016; Kowalska-Krochmal & Dudek-Wicher, 2021). For turbidity-based quantification, the untreated bacterial growth control, blank control, and sample background control were included in parallel. The blank control contained sterile medium only, whereas the sample background wells contained antibiotic-free medium supplemented with the corresponding concentration of DAAT preparation but no bacteria, allowing correction for any intrinsic turbidity caused by the treatment sample(Balouiri et al., 2016; Kowalska-Krochmal & Dudek-Wicher, 2021) . Each condition was tested in at least three parallel replicates. The bacterial growth inhibition rate was calculated from background-corrected *OD₆₀₀* values according to the following equation:

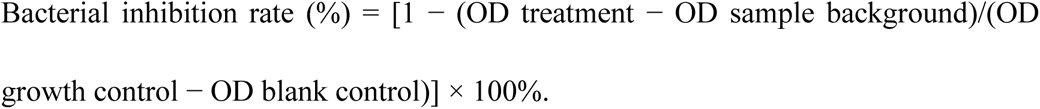

In this equation, OD treatment represents the *OD₆₀₀* value obtained from bacteria incubated with the indicated DAAT preparation; OD sample background represents the *OD₆₀₀*value of medium containing the corresponding DAAT preparation without bacteria; OD growth control represents the *OD₆₀₀* value of untreated bacteria grown under the same conditions; and OD blank control represents the *OD₆₀₀* value of sterile medium alone. The inhibitory effect of each DAAT formulation against *Staphylococcus aureus* was determined by comparing the corrected turbidity of treated wells with that of the untreated bacterial growth control.

### 2.12 Quantification of intracellular *Staphylococcus aureus* burden by host-cell lysis and spot plating

To evaluate the antibacterial activity of DAAT formulations against intracellular *Staphylococcus aureus*, infected HT-29 cells were treated with DAAT Cecropin, DAAT LL37, or DAAT Indolicidin at the optimized concentrations of 256 and 512 ng μL⁻¹. After 12 h of treatment under standard culture conditions, the culture medium was removed, and the cells were gently washed with sterile PBS to eliminate residual extracellular components. Cells were then detached by digestion with trypsin at 37 °C for 4 min, and an equal volume of complete medium was added to terminate enzymatic digestion. The cell suspension was transferred into 1.5 mL sterile centrifuge tubes and collected for intracellular bacterial recovery. To release intracellular bacteria, the collected cells were resuspended in lysis buffer containing 1% Triton X-100 and incubated on ice for 30 min to ensure sufficient disruption of host-cell membranes while preserving bacterial viability (Häffner et al., 2020; Peyrusson et al., 2022). The lysates were centrifuged at 12,000 × g for 10 min at 4 °C, and the resulting supernatants were collected for bacterial burden analysis. Serial ten-fold dilutions of each lysate were prepared in sterile PBS or TSB medium up to 10⁻⁵. For each dilution, 5 μL of diluted lysate was spotted onto TSB agar plates. After the droplets were fully absorbed into the agar surface, the plates were inverted and incubated overnight at 37 °C for 10 h. Colony growth was then photographed and recorded (Häffner et al., 2020). Untreated infected cells and normal uninfected cells were included as controls. The reduction in colony formation after DAAT treatment was used to assess the clearance efficiency of each formulation against intracellular *S. aureus*.

### 2.13 Safety evaluation of DAAT formulations in HT-29 cells

To evaluate the Safety evaluation of DAAT-Cecropin, DAAT-LL37, and DAAT-Indolicidin toward host cells, HT-29 cells were seeded into 96-well cell culture plates in complete culture medium and allowed to attach and grow uniformly under standard culture conditions. After stable cell adherence, the medium was removed and replaced with fresh complete medium containing serial concentrations of DAAT-Cecropin, DAAT-LL37, or DAAT-Indolicidin. For each formulation, the final concentrations were set at 512, 256, 128, 64, 32, 16, 8, 4, 2, and 1 ng μL⁻¹. Untreated HT-29 cells cultured in complete medium were used as control group, whereas medium-only wells were included as blank controls to correct background signal. After 24 h of incubation, HT-29 cell viability was assessed using a cell viability assay kit HT-29 cell viability was measured using PrestoBlue™ Cell Viability Reagent (Thermo Fisher Scientific, Waltham, MA, USA; Cat. No. A13261) according to the manufacturer’s instructions (Thermo Fisher Scientific, 2024). Briefly, PrestoBlue reagent was added directly to each well at the recommended volume, and the plates were incubated at 37 °C until the signal reached the linear detection range. Fluorescence was then measured using a microplate reader at 560/590 nm excitation/emission. When absorbance detection was used, readings were collected at 570 nm with 600 nm as the reference wavelength. The relative growth rate of HT-29 cells was calculated by normalizing the signal from treated wells to that of the untreated control after blank subtraction, following the calculation method provided by the manufacturer. Each treatment condition was tested in triplicate, and the resulting results were used to determine the concentration range in which DAAT formulations showed acceptable compatibility with HT-29 cells. In parallel, representative HT-29 cells treated with DAAT-LL37 at 512, 256, 128, 64, and 32 ng μL^-1^ were imaged by pseudocolor optical microscopy, following the imaging procedure and parameter settings described in the section “ Pseudocolor optical microscopy imaging.”

### 2.14 Live-cell fluorescence imaging of HT-29 cells

HT-29 cell viability and growth after repeated DAAT treatment were assessed by live/dead fluorescence staining. Infected HT-29 cells were divided into four groups: untreated infected control, DAAT-Cecropin, DAAT-LL37, and DAAT-Indolicidin. Treatment groups received the corresponding DAAT formulation at 256 ng μL⁻¹, with dosing repeated every 24 h. Cell status was examined after one, two, and three rounds of treatment, while the control group received an equal volume of fresh medium at the same time points. At each time point, the medium was removed, and cells were gently washed with sterile PBS to remove residual medium and unbound DAAT particles. A live/dead staining solution containing Calcein-AM and propidium iodide (MCE, HY-K1094) was prepared at final concentrations of 2 μM and 4.5 μM, respectively, and added to the cells (MedChemExpress, 2024). Viable cells were identified by green fluorescence from Calcein-AM staining, whereas dead cells were labeled by propidium iodide (Suzuki et al., 2017). Because detached dead cells may be lost during washing, the green fluorescence channel was primarily used to evaluate surviving adherent cells. After staining, cells were gently rinsed with PBS and imaged immediately under a fluorescence microscope. Representative images were acquired using a 4× objective lens with the exposure intensity set to 60%. Identical imaging parameters were applied across all groups and time points. Green fluorescence-positive cells were counted from each image using an AI-assisted image analysis workflow, and the cell counts were used to compare live-cell abundance after repeated DAAT treatment (O’Brien et al., 2016).

### 2.15 Inflammatory cytokine analysis after DAAT treatment

To evaluate the inflammatory response of infected HT-29 cells following DAAT treatment, the concentrations of tumor necrosis factor-α (TNF-α), interleukin-1β (IL-1β), interleukin-6 (IL-6) and interferon-γ (IFN-γ) were measured after a single administration (Deramaudt et al., 2020; Liu et al., 2021; Luqman et al., 2025). Intracellular *Staphylococcus aureus*-infected HT-29 cells were assigned to an untreated infection control group and DAAT-Cecropin, DAAT-LL37 and DAAT-Indolicidin treatment groups. Each treatment group received the corresponding DAAT formulation at the designated working concentration, whereas the control group was treated with an equal volume of PBS. At the indicated endpoint, cell-culture supernatants were collected and centrifuged to remove detached cells and particulate debris. The clarified supernatants were stored at −80 °C until analysis. TNF-α, IL-1β, IL-6 and IFN-γ concentrations were quantified using human ELISA kits according to the manufacturer’s instructions (Solarbio, Beijing, China; Cat. Nos. SEKH-0047, SEKH-0002, SEKH-0013 and SEKH-0046, respectively) (Liu et al., 2021). Cytokine concentrations were calculated by interpolation from standard curves generated in parallel. All experimental groups were analyzed using three independent replicates.

### 2.16 Direct antibacterial activity of tail-fibre-retargeted DAAT

The amino acid sequences of the original and modified tail-fibre proteins were submitted to AlphaFold2 for structural prediction. Five models were generated for each protein sequence, and the highest-ranked model was selected according to the predicted local distance difference test score. The predicted structures were visualized in PyMOL to compare the overall architecture of the original and engineered tail-fibre proteins.

The direct antibacterial activity of tail-fibre-retargeted DAAT against extracellular *Staphylococcus aureus* was evaluated by a colony-forming assay (Clinical and Laboratory Standards Institute, 2024). A bacterial suspension with an OD₆₀₀ of 1.0, corresponding to approximately 5 × 10⁷ cells mL⁻¹, was prepared and diluted as required. For each reaction, 1.28 μL of the bacterial suspension, equivalent to 6.4 × 10⁴ CFU, was mixed with 50, 100 or 200 μL of tail-fibre-retargeted DAAT at 600 ng μL⁻¹. The control group was treated with an equal volume of PBS under identical conditions. The mixtures were incubated for 4 h under the indicated culture conditions. After incubation, each sample was serially diluted by 10-fold dilution to 10⁻². Subsequently, 100 μL of each dilution was spread onto TSB agar plates and incubated under standard bacterial culture conditions. Colonies were imaged and counted after incubation, and the viable bacterial burden was expressed as CFU mL⁻¹. The reduction in colony formation relative to the untreated control was used to evaluate the direct antibacterial activity of tail-fibre-retargeted DAAT against extracellular *S. aureus*.

### 2.17 Mouse subcutaneous Staphylococcus aureus infection and DAAT treatment

Wild-type C57BL/6 mice were bred and maintained in-house under standard laboratory conditions. Age-appropriate mice at a normal stage of growth and development were selected for the experiments. A mouse subcutaneous *Staphylococcus aureus* infection model was established by injecting 100 μL of bacterial suspension into the dorsal subcutaneous tissue (Kim et al., 2014; Malachowa et al., 2019; Subramanian et al., 2021). The bacterial suspension was adjusted to OD₆₀₀ = 1.0, corresponding to approximately 5 × 10⁷ cells mL⁻¹; thus, each mouse received approximately 5 × 10⁶ bacterial cells. Infection was allowed to develop for 2 days, after which visible signs of subcutaneous infection were confirmed before treatment. Mice were then treated locally with different DAAT formulations designed for coordinated intracellular and extracellular bacterial clearance. Each 100 μL dose contained 50 μL of intracellular AMP-delivering DAAT at 256 ng μL⁻¹ and 50 μL of tail-fibre-retargeted DAAT directed against extracellular *S. aureus* at 600 ng μL⁻¹, corresponding to 12.8 μg and 30 μg of the respective DAAT formulations per administration. Treatment was administered once every 24 h for 6 days. Wounds of representative mice from each group were photographed every 72 h (Malachowa et al., 2019; Subramanian et al., 2021).

### 2.18 Histological analysis of mouse wound tissues by H&E and Masson ’ s trichrome staining

Histological examination was conducted to assess wound repair, inflammatory infiltration, tissue regeneration, and collagen deposition in mouse wound tissues. On day 12 after treatment, the wound area together with adjacent normal skin was carefully excised and fixed in 4% paraformaldehyde for 24–48 h. The fixed specimens were dehydrated through graded ethanol, cleared with xylene, embedded in paraffin, and sectioned into 4–5 μm slices. For hematoxylin and eosin (H&E) staining, paraffin sections were deparaffinized, rehydrated, stained with hematoxylin, differentiated, blued, counterstained with eosin, dehydrated, cleared, and mounted with neutral resin. The H&E-stained sections were examined under a light microscope to evaluate re-epithelialization, granulation tissue formation, inflammatory cell infiltration, and overall tissue morphology. For Masson’s trichrome staining, deparaffinized and rehydrated sections were stained using a Masson’s trichrome staining kit according to the manufacturer’s instructions (Solarbio, Beijing, China; Cat. No. G1340). Collagen fibers appeared blue, whereas muscle fibers and cytoplasm appeared red. The stained sections were used to assess collagen accumulation, fiber organization, and extracellular matrix remodeling during wound healing. Images were captured under a light microscope, and histological differences among groups were compared based on inflammation, epidermal regeneration, granulation tissue formation, and collagen deposition.

### 2.19 Statistics and reproducibility

Statistical analyses were performed using IBM SPSS Statistics 23. Quantitative data are shown as mean ± SD from at least three independent experiments unless otherwise stated. Graphs and schematic figures were prepared with Origin 2024 and Adobe Illustrator 2020. Comparisons among multiple groups were conducted using one-way or two-way ANOVA followed by Bonferroni correction. Statistical significance was defined as *P* < 0.05, with significance levels indicated as *P* < 0.05 (*), *P* < 0.01 (**), and *P* < 0.001 (***).

## 3. Result

### 3.1 Engineering and validation of antimicrobial peptide-loaded DAAT particles

To assess the antibacterial activity of the selected peptides, Cecropin, LL-37, and Indolicidin were chemically synthesized and tested against *Staphylococcus aureus* using a concentration-gradient plate assay (Figure 1B). At 32–128 ng μL⁻¹, all three peptides showed limited bacterial clearance, as indicated by dense colony formation. When the concentration increased to 256 and 512 ng μL⁻¹, colony numbers were markedly reduced, demonstrating effective killing of *S. aureus* at higher doses. This dose-dependent activity is consistent with the membrane-targeting behavior of cationic antimicrobial peptides. After binding to the negatively charged bacterial surface, these peptides disrupt membrane integrity through different modes: Cecropin mainly forms membrane pores, LL-37 induces carpet-like membrane permeabilization, and Indolicidin destabilizes membranes and can further interact with intracellular targets. These actions promote ion leakage, cytoplasmic content release, and loss of bacterial viability (Figure 1A) (Silvestro et al., 2000; Lee et al., 2011; Hsu et al., 2005; Le et al., 2017). The results confirm that Cecropin, LL-37, and Indolicidin possess effective anti-*S. aureus* activity at sufficient concentrations, supporting their use as functional payloads for DAAT-mediated intracellular antibacterial delivery.

**Figure 1.**
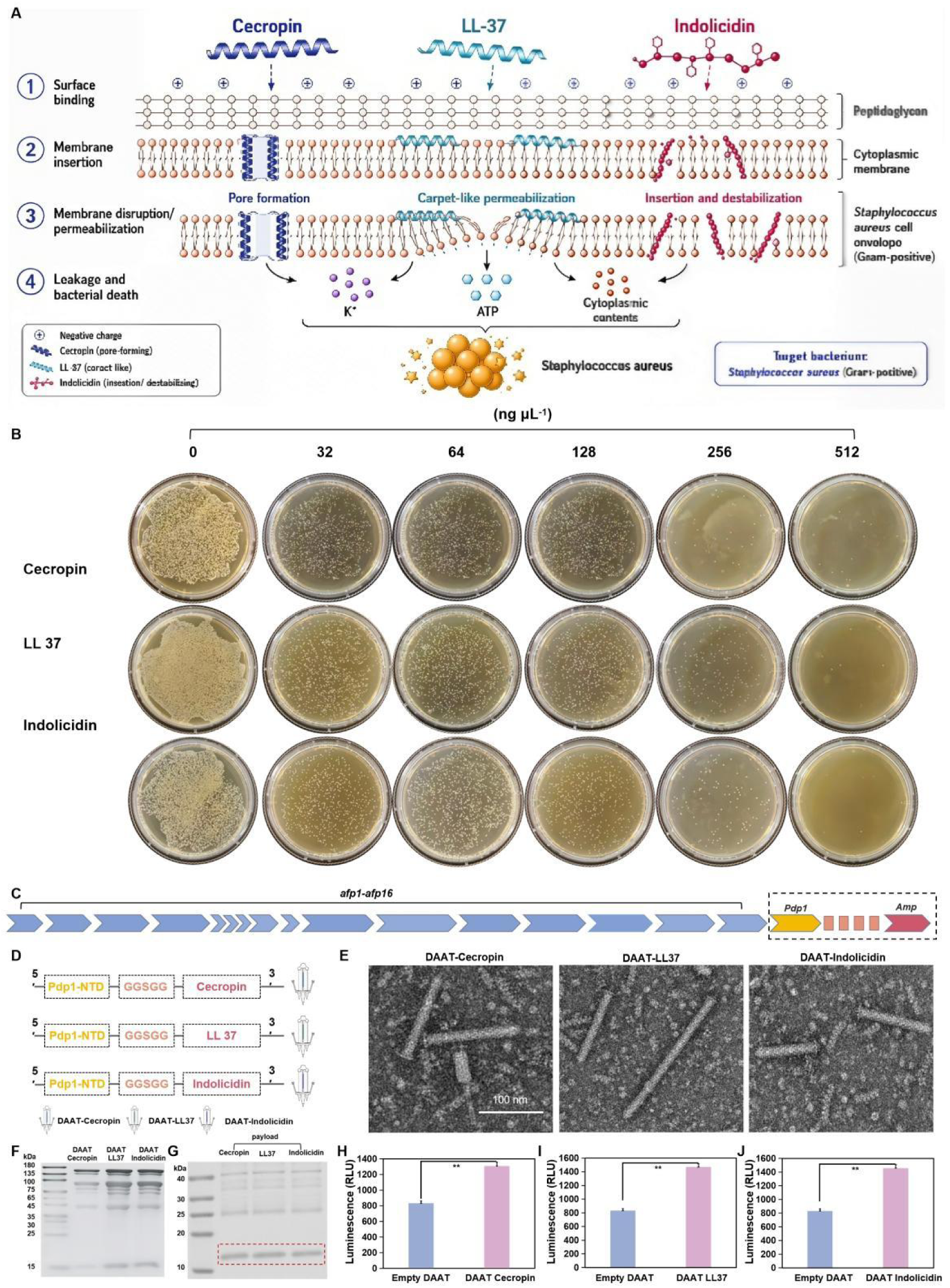
Design, construction and validation of antimicrobial peptide-loaded DAAT particles. (A) Schematic illustration of the antibacterial mechanisms of Cecropin, LL-37 and Indolicidin against *Staphylococcus aureus*. Cecropin mainly induces pore formation, LL-37 promotes carpet-like membrane permeabilization, and Indolicidin inserts into and destabilizes the bacterial membrane, ultimately causing ion leakage, cytoplasmic content release and bacterial death. (B) Concentration-gradient plate assay showing the antibacterial activities of synthetic Cecropin, LL-37 and Indolicidin against *S. aureus*. Peptides were tested at 0, 32, 64, 128, 256 and 512 ng μL⁻¹, and colony formation decreased with increasing peptide concentration. (C) Genetic organization of the DAAT system. The *afp1–afp16* structural gene cluster encodes the contractile injection apparatus, while the payload cassette contains the Pdp1-derived loading element and antimicrobial peptide cargo. (D) Construction of DAAT-Cecropin, DAAT-LL37 and DAAT-Indolicidin payloads by fusing each antimicrobial peptide downstream of Pdp1-NTD through a flexible GGSGG linker. (E) Negative-stain TEM images of purified DAAT-Cecropin, DAAT-LL37 and DAAT-Indolicidin particles showing rod-shaped PVC-like nanosyringe morphology. Scale bar, 100 nm. (F) SDS-PAGE analysis of purified DAAT particles showing comparable high-molecular-weight structural protein profiles among different DAAT variants. (G) Low-molecular-weight gel analysis showing specific antimicrobial peptide-containing fusion bands in the Cecropin, LL37 and Indolicidin lanes, confirming the expression of peptide payloads. (H–J) HiBiT-based luminescence assays confirming increased payload signals in DAAT-Cecropin, DAAT-LL37 and DAAT-Indolicidin particles compared with empty DAAT, respectively. Data are presented as mean ± SD; \*\**P < 0.01*.

To achieve the safe expression and protective delivery of AMPs, we engineered the loading elements of PVC pnf nanosyringes via fusion pdp1 and AMPs to construct a versatile DAAT delivery system capable of carrying various replaceable peptide payloads (Vlisidou et al., 2019; Jiang et al., 2022; Kreitz et al., 2023). This work extends the application of the PVC system to therapeutic intervention against intracellular bacteria, providing a new strategy for intracellular bacterial research. The backbone of DAAT consists of the pvc1–pvc16 structural elements encoding the contractile injection system, the Pdp1-derived leading sequence for payload loading, and payload elements carrying antimicrobial peptides (Amps) against Staphylococcus aureus(Figure 1C) (Jiang et al., 2019; Vlisidou et al., 2019; Jiang et al., 2022; Kreitz et al., 2023). This design enables co-expression of structural particles and AMP payload modules in Escherichia coli, allowing particle assembly and encapsulation of AMP payloads to be accomplished within a single host strain (Jiang et al., 2022; Kreitz et al., 2023).

To confirm successful intraluminal loading of antimicrobial peptides into PVC particles, three representative AMPs, including Cecropin, LL-37, and Indolicidin, were selected as model cargos. We design a sophisticated strategy for loading AMPs, where the AMP was fused with PD via a flexible GGSGG linker peptide (Figure 1D). In this design, pdp1 guide AMP loaded into PVC inner tube, whereas the glycine-rich flexible linker enables to reduce steric interference between the packaging module and the antimicrobial peptide sequence (Chen et al., 2013; Jiang et al., 2022). Based on this strategy, three DAAT variants were constructed and designated DAAT-Cecropin, DAAT-LL37 and DAAT-Indolicidin. To determine whether AMP loading affected particle assembly, we examined purified DAAT particles by negative-stain transmission electron microscopy. All three preparations contained rod-shaped nanosyringe-like particles with PVC-like morphology, including an elongated sheath–tube body and a terminal baseplate-associated region, consistent with assembled extracellular contractile injection particles. (Figure 1E) (Jiang et al., 2019; Desfosses et al., 2019). These results further suggest that the PVC structural module is compatible with the modified peptide-loading cassette. SDS-PAGE analysis further showed that the major high-molecular-weight bands in all three DAAT preparations corresponded to the principal structural proteins of PVC. (Figure 1F) (Jiang et al., 2019; Kreitz et al., 2023). No marked differences were detected in the electrophoretic profiles, indicating that all three recombinant systems generated PVC-derived particles with comparable structural protein compositions. Thus, the introduction of distinct antimicrobial peptide cargos did not substantially alter the overall assembly characteristics or structural protein composition of DAAT particles. After confirming intact particle assembly, we further validated whether AMP payloads were efficiently incorporated into purified DAAT particles. A HiBiT luminescence assay was applied for detection of peptide cargos. The HiBiT tag was fused to the C-terminus of AMP sequences, and analysis of AMP loading was achieved through high-affinity complementation between HiBiT and LgBiT subunits (Schwinn et al., 2018; Schwinn et al., 2020). Compared with the empty DAAT control group, all three AMP-loaded DAAT particles exhibited markedly enhanced luminescence signals (Figure 1G). The luminescence intensities of DAAT-LL37 and DAAT-indolicidin were slightly higher than that of DAAT-cecropin, indicating that the engineered peptide payloads were stably retained inside DAAT particles and did not suffer substantial loss during purification. Furthermore, protein gel electrophoresis was used to verify the presence of low-molecular-weight AMP payloads. Distinct specific bands were detected at the molecular weight region corresponding to target cargos in the lanes of cecropin, LL-37 and indolicidin. All bands were approximately 11 kDa, which matched the theoretical molecular weight of the fusion protein composed of AMP and HiBiT tag (Figure 1H-J). This provides direct evidence for successful payload of AMP.

Collectively, the results from TEM morphological observation, SDS-PAGE and HiBiT luminescence assay demonstrate that cecropin, LL-37 and indolicidin were successfully loaded into DAAT inner tube. In this study, a panel of AMP-loaded nanosyringes was successfully constructed, laying a solid foundation for subsequent evaluation of antibacterial activity.

### 3.2 Antimicrobial DAAT treatments intracellular *S. aureus*

After confirming that antimicrobial peptide cargos could be loaded into the DAAT lumen,, we next examined whether DAAT loaded with AMPs could functionally suppress intracellular S. aureus infection. To establish a tractable infection model, HT-29 cells were infected with the GFP-expressing USA300/Eno-GFP strain, enabling bacterial colonization to be monitored by fluorescence imaging (Luqman et al., 2019; Bravo-Santano et al., 2019). Uninfected HT-29 cells maintained a typical adherent epithelial morphology, whereas infected cells exhibited clear green fluorescence together with infection-associated morphological disturbance (Figure 2A) (Rodrigues Lopes et al., 2022). To define a suitable bacterial input for subsequent treatment assays, we compared extracellular and intracellular bacterial burdens across increasing inocula. As expected, extracellular bacterial counts increased sharply with increasing bacterial inocula, whereas the intracellular bacterial burden gradually reached a plateau. Considering both detectable intracellular colonization and preservation of host-cell status, an inoculum of 6.4 × 10⁴ CFU per well was selected as the optimal infection dose for subsequent experiments (Figure 2B).

**Figure 2.**
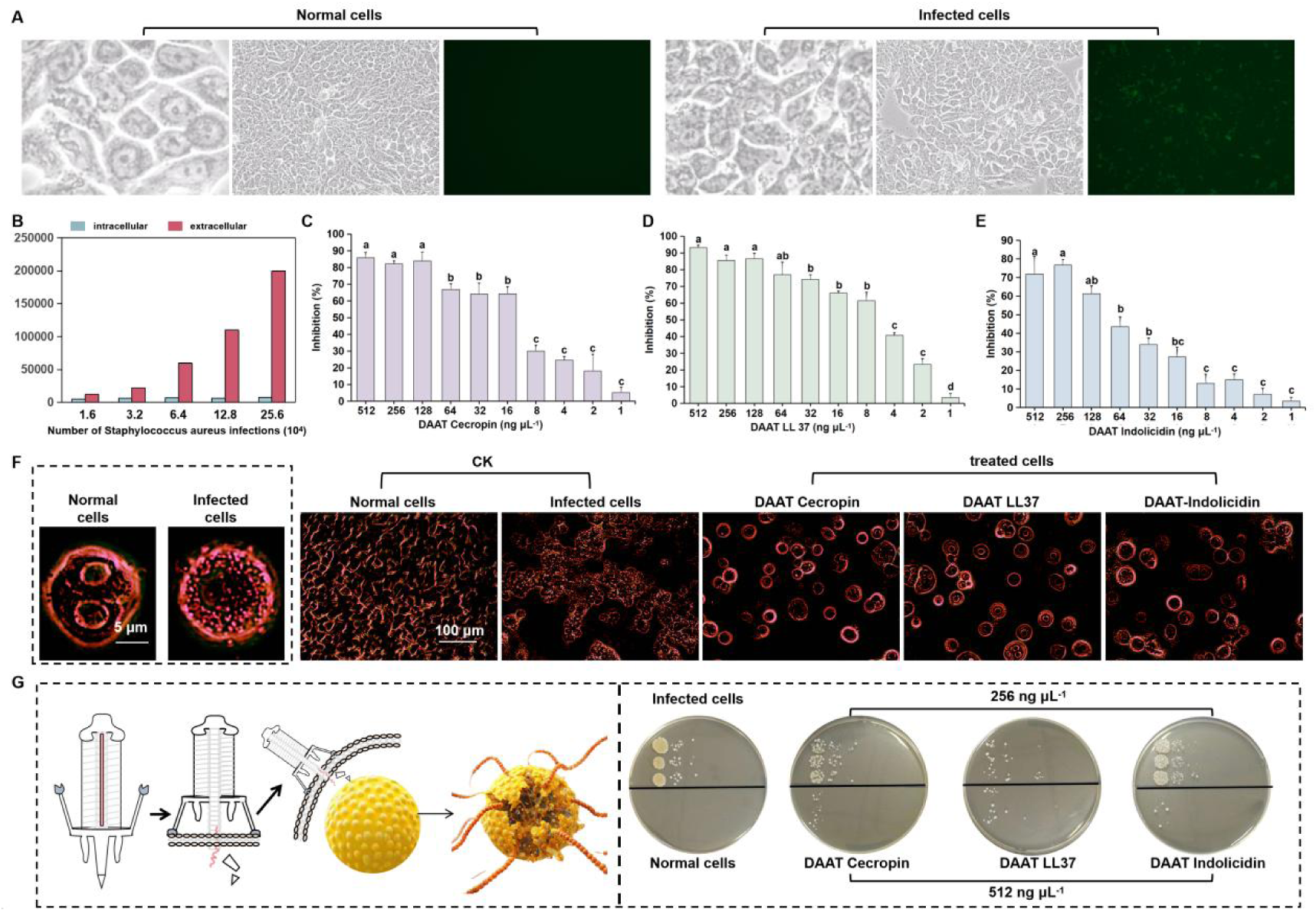
DAAT-mediated antimicrobial peptide delivery suppresses intracellular *Staphylococcus aureus* infection in HT-29 cells. (A) Bright-field and GFP fluorescence images of normal HT-29 cells and USA300/Eno-GFP-infected HT-29 cells, confirming successful establishment of the intracellular *S. aureus* infection model. (B) Intracellular and extracellular bacterial burdens in HT-29 cells infected with different bacterial inocula. The intracellular bacterial load increased with infection dose, and 6.4 × 10⁴ CFU per well was selected for subsequent experiments based on efficient intracellular infection with relatively low extracellular bacterial contamination. (C–E) Dose-dependent inhibition of intracellular *S. aureus* by DAAT-Cecropin, DAAT-LL37, and DAAT-Indolicidin, respectively. (F) Pseudocolor microscopy images showing the morphological changes of normal, infected, and DAAT-treated HT-29 cells. Infection caused obvious cell shrinkage, aggregation, and structural disorder, whereas DAAT-Cecropin, DAAT-LL37, and DAAT-Indolicidin treatments partially reduced infection-associated cellular damage. Scale bars, 5 μm and 100 μm, as indicated. (G) Schematic illustration of DAAT-mediated antimicrobial peptide delivery into infected cells and colony spot-plating assay of recoverable intracellular bacteria after treatment. Compared with infected cells, DAAT-Cecropin, DAAT-LL37, and DAAT-Indolicidin markedly reduced intracellular bacterial survival at 256 and 512 ng μL⁻¹. Data are presented as mean ± SD. Different lowercase letters indicate significant differences among treatments (*P < 0.05*)

Having established the infection window, we then evaluated the antibacterial activity of three peptide-loaded DAAT formulations after repeated dosing. DAAT Cecropin, DAAT LL37 and DAAT Indolicidin were applied over a concentration range from 1 to 512 ng μL⁻¹, and bacterial inhibition was assessed after three rounds of treatment. All three formulations showed dose-dependent suppression of bacterial growth, but their activity profiles differed. DAAT Cecropin maintained a high inhibitory effect across the upper concentration range, with strong inhibition still observed at intermediate doses (Figure 2C). DAAT LL37 also produced pronounced antibacterial activity, showing a gradual decline as the dose decreased, consistent with concentration-dependent intracellular bacterial control (Figure 2D). DAAT Indolicidin displayed a narrower active window, with robust inhibition at higher concentrations but a steeper loss of efficacy at lower doses (Figure 2E). These results indicate that the antibacterial output was shaped by the identity and potency of the loaded antimicrobial peptide.

Pseudocolor optical imaging distinguished normal cells from infected cells and provided a direct visual readout of cellular status after treatment. Normal HT-29 cells formed a continuous adherent layer with regular cellular outlines, whereas infected cells showed marked morphological deterioration (Figure 2F) (Rodrigues Lopes et al., 2022). After of DAAT treatment, the infected-cell phenotype was partially reversed, with treated groups displaying fewer severely damaged cellular regions than the untreated infected control. Thus, the microscopy data supported the view that DAAT-mediated delivery reduced infection-associated cellular injury rather than merely altering turbidity-based readouts.

To directly assess intracellular bacterial colonization after treatment, we next performed host-cell lysis followed by serial-dilution spot plating (Figure 2G) (Bravo-Santano et al., 2019). Infected cells produced visible colony growth after plating, whereas normal cells showed no colony formation. After treatment of infected HT-29 cells with DAAT particles, the number of intracellular colonies was markedly reduced. Taking the treatment doses of 256 ng μL⁻¹ and 512 ng μL⁻¹ as examples, all DAAT formulations showed antibacterial activity, with DAAT-LL37 exhibiting the strongest therapeutic effect against intracellular bacteria, followed by DAAT-Cecropin and DAAT-Indolicidin. In addition, DAAT particle treatment significantly decreased the colony-forming ability of infected HT-29 cells, indicating that DAAT therapy displayed a concentration-dependent antibacterial trend. More importantly, the treatment dose that can maintain normal host-cell growth while controlling intracellular bacterial infection needs to be further defined.

### 3.3 Dose optimization of DAAT treatment

Having established that DAAT particles could deliver antimicrobial peptide cargos and suppress intracellular *Staphylococcus aureus*, we next asked whether this antibacterial activity could be achieved within a concentration window compatible with HT-29 host-cell growth. This issue was essential because an effective intracellular antibacterial platform should not simply reduce bacterial burden, but should also preserve the recovery potential of infected host cells (Greco et al., 2020; Bellavita et al., 2023). To define this working range, normal HT-29 cells were exposed to DAAT Cecropin, DAAT LL37 and DAAT Indolicidin across a concentration gradient from 1 to 512 ng μL⁻¹ (Figure 3A-C). In contrast, 512 ng μL⁻¹ consistently caused a visible reduction in HT-29 cell growth, indicating that this highest dose entered a host-cell stress range rather than an optimal therapeutic window. Thus, although 512 ng μL⁻¹ retained antibacterial potential, it was not selected as the preferred treatment dose.

**Figure 3.**
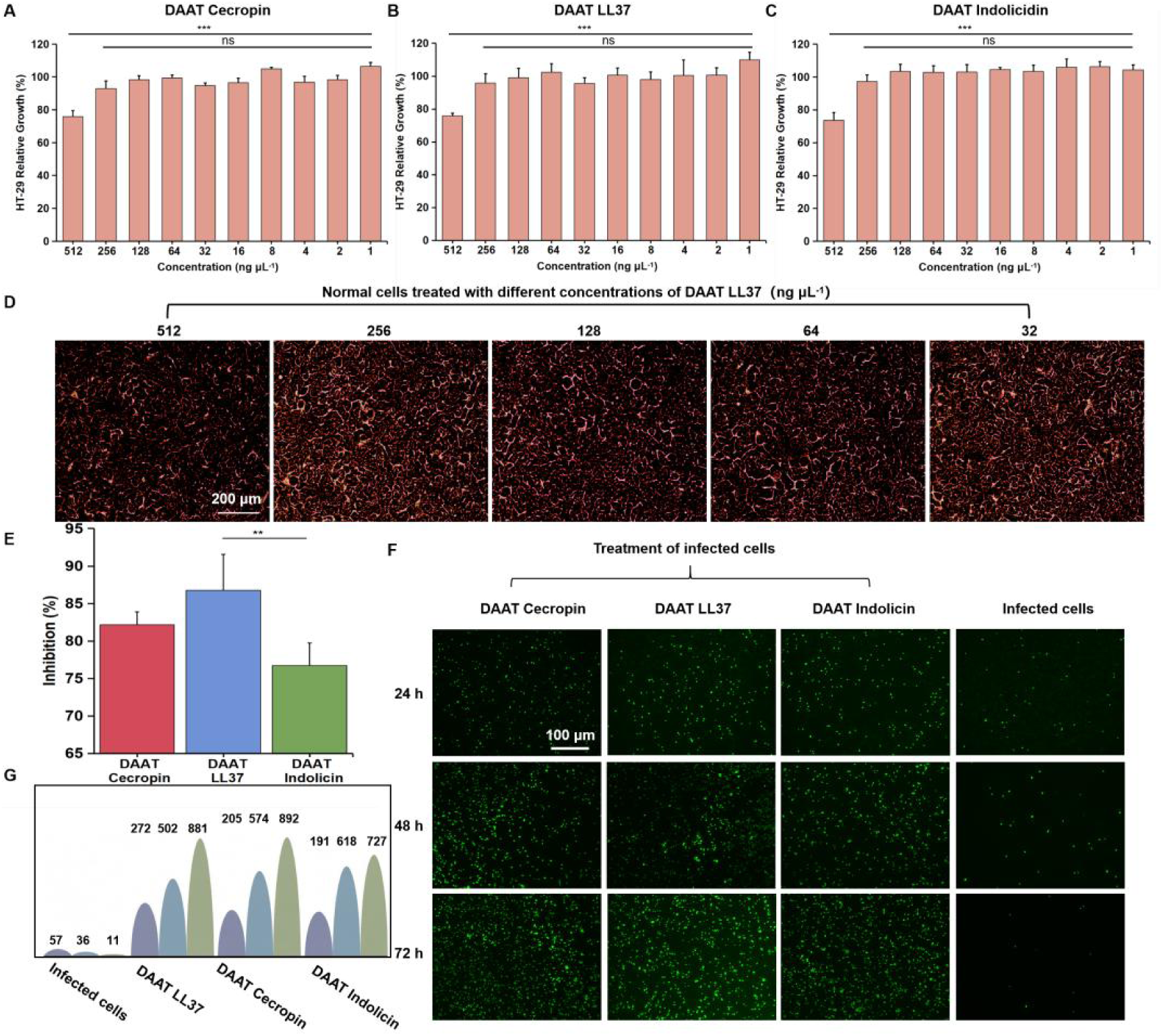
Cytocompatibility and therapeutic efficacy of DAAT treatment. (A–C) Cytocompatibility of DAAT-Cecropin, DAAT-LL37 and DAAT-Indolicidin in normal HT-29 cells. All formulations were well tolerated across most tested concentrations, whereas 512 ng μL⁻¹ reduced cell growth. Data are shown as mean ± SD; \**P* < 0.05, \*\**P* < 0.01; ns, not significant. (D) Pseudocolor optical images of HT-29 cells treated with different concentrations of DAAT-LL37. Cells maintained favourable morphology at 256 ng μL⁻¹ and lower concentrations. Scale bar, 200 μm. (E) Antibacterial inhibition of three DAAT formulations at 256 ng μL⁻¹, with DAAT-LL37 showing the strongest effect. Data are shown as mean ± SD; \**P* < 0.01. (F) Live-cell fluorescence images of infected HT-29 cells after repeated DAAT treatment at 256 ng μL⁻¹ for 24, 48 and 72 h. Green fluorescence indicates surviving adherent cells. Scale bar, 100 μm. (G) Quantification of fluorescence-positive live cells, showing progressive recovery after repeated DAAT treatment.

This concentration-dependent safety profile was further supported by pseudocolor optical microscopy using DAAT LL37 as a representative formulation. At 256, 128, 64 and 32 ng μL⁻¹, HT-29 cells retained a dense and continuous adherent morphology, with preserved cellular spreading and organized growth patterns. By comparison, cells exposed to 512 ng μL⁻¹ showed a relatively less favorable morphology, consistent with the quantitative viability data. These imaging results provided a visual confirmation that the decrease observed at the highest dose reflected genuine cellular stress rather than only assay fluctuation (Figure D). Based on the combined viability and morphology evidence, 256 ng μL⁻¹ was selected as a more suitable working concentration for subsequent intracellular infection treatment, as it retained strong antibacterial capacity while avoiding the cytotoxicity associated with the 512 ng μL⁻¹ condition.

To verify the therapeutic efficacy of repeated dosing at the optimal treatment concentration, we examined whether repeated DAAT treatment could progressively improve the growth status of infected cells over time. Live/dead fluorescence staining revealed that untreated infected cells maintained only sparse green fluorescence, reflecting poor survival and limited adherent-cell recovery after infection. In contrast, cells treated with DAAT Cecropin, DAAT LL37 or DAAT Indolicidin showed progressively increased green fluorescence from 24 to 72 h (Figure 3F and G). The repeated-dosing results showed that infected HT-29 cells gradually regained normal proliferative capacity during successive DAAT treatments. Therefore, all DAAT-treated groups showed an increase in live-cell numbers over successive treatment cycles, whereas the infected control remained at a much lower level. Among the three formulations, DAAT LL37 produced the most pronounced recovery trend, in agreement with its higher bacterial inhibition rate at 256 ng μL⁻¹.

Together, these results define a functional treatment window for DAAT-mediated intracellular antibacterial therapy. Therefore, 256 ng μL⁻¹ was selected as the optimal concentration for antibacterial treatment. Under this condition, repeated DAAT peptide treatment reduced intracellular bacterial pressure and allowed infected HT-29 cells to progressively recover their adherent growth state. Therefore, DAAT Cecropin, DAAT LL37 and DAAT Indolicidin do not only act as antibacterial particles against intracellular *S. aureus*; within an appropriate dose range, they also preserve host-cell survival.

### 3.4 DAAT reduces inflammatory cytokine levels in infected cells

Having shown that DAAT particles suppressed intracellular Staphylococcus aureus, we next examined whether this antibacterial effect was accompanied by reduced inflammatory cytokine responses in infected HT-29 cells. Four representative inflammatory cytokines, including TNF-α, IL-1β, IFN-γ and IL-6, were quantified after treatment with DAAT Cecropin, DAAT LL37 and DAAT Indolicidin (Figure 4A-D). Compared with the infected control group, all DAAT-treated groups showed a clear decrease in cytokine levels, indicating that DAAT-mediated intracellular bacterial inhibition was associated with attenuation of the infection-induced inflammatory response.

**Figure 4.**
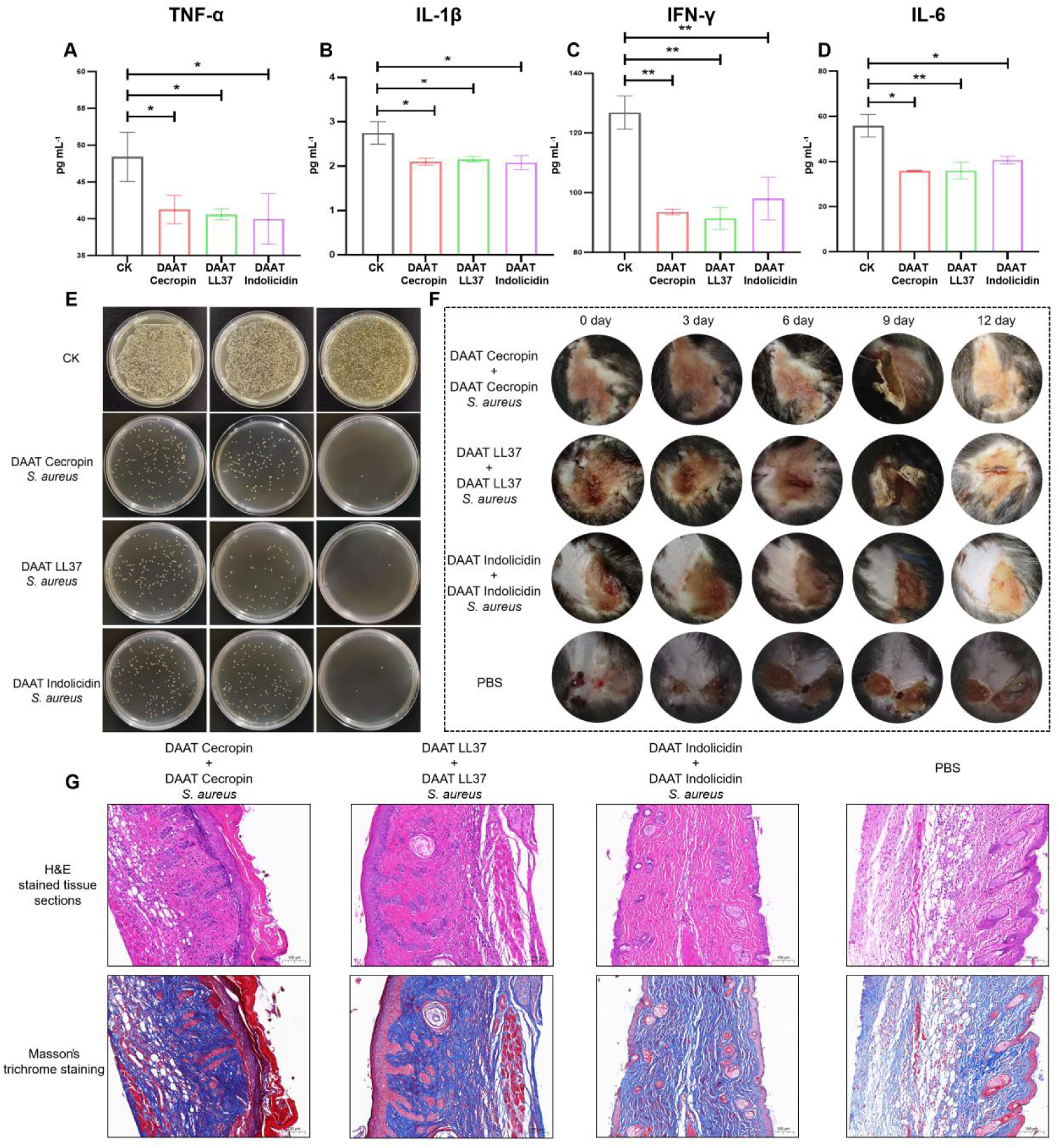
DAAT-mediated antimicrobial peptide delivery reduces inflammatory responses, kills retargeted *Staphylococcus aureus*, and promotes skin wound healing in mice. (A–D) Quantification of inflammatory cytokines after DAAT treatment, including TNF-α, IL-1β, IFN-γ, and IL-6. Compared with the infected control group, DAAT-Cecropin, DAAT-LL37, and DAAT-Indolicidin reduced cytokine levels, indicating attenuation of the infection-induced inflammatory response following intracellular *S. aureus* inhibition. (E) Colony-forming assay showing the direct antibacterial activity of tail-fibre-retargeted DAAT against extracellular *S. aureus*. Bacteria were incubated with DAAT-Cecropin-*S. aureus*, DAAT-LL37-*S. aureus*, or DAAT-Indolicidin-*S. aureus*, followed by serial dilution and plating on TSB agar. Compared with the CK group, all retargeted DAAT formulations markedly reduced colony formation, confirming effective extracellular bacterial killing after tail-fibre modification. (F) Representative images of infected skin wounds in a murine subcutaneous *S. aureus* infection model during treatment. Wounds were recorded on days 0, 3, 6, 9, and 12 after treatment with DAAT-Cecropin-*S. aureus*, DAAT-LL37-*S. aureus*, DAAT-Indolicidin-*S. aureus*, or PBS. DAAT-treated groups showed reduced wound exudation, faster scab formation, and improved wound closure compared with PBS-treated mice. (G) Histological analysis of infected skin tissues by hematoxylin and eosin (H&E) staining and Masson’s trichrome staining. DAAT-treated groups exhibited reduced inflammatory infiltration, improved epidermal organization, and enhanced collagen deposition compared with the PBS group, indicating accelerated tissue repair and remodeling. Scale bars, 100 μm. Data are presented as mean ± SD. \**P < 0.05*, \*\**P < 0.01*.

For TNF-α, the control group showed a level of approximately 48.3, whereas DAAT Cecropin, DAAT LL37 and DAAT Indolicidin reduced the level to about 41.2, 40.6 and 40.0, respectively. A similar trend was observed for IL-1β, which decreased from approximately 2.7 in the control group to 2.1, 2.1 and 2.0 after DAAT Cecropin, DAAT LL37 and DAAT Indolicidin treatment, respectively. IFN-γ showed the most pronounced reduction among the tested cytokines. Its level declined from approximately 126.5 in the control group to 93.0, 91.5 and 98.0 in the three DAAT treatment groups, respectively. In addition, IL-6 was reduced from approximately 55.0 in the control group to 35.5, 35.0 and 40.5 after treatment with DAAT Cecropin, DAAT LL37 and DAAT Indolicidin, respectively.

Statistical analysis further confirmed that DAAT treatment significantly reduced the levels of these inflammatory cytokines compared with the infected control group. Overall, these results demonstrate that DAAT Cecropin, DAAT LL37 and DAAT Indolicidin not only inhibit intracellular S. aureus, but also alleviate the inflammatory cytokine response induced by intracellular infection.

### 3.5 Tail-fiber modification enables DAAT-mediated targeted killing of Staphylococcus aureus

To examine whether tail-fibre retargeting was compatible with DAAT assembly and bacterial targeting, the direct antibacterial activity of tail-fibre-retargeted DAAT against extracellular S. aureus was then evaluated by colony-forming assay (Figure 4E). An initial bacterial input of 6.4 × 10⁴ CFU was incubated with DAAT Cecropin S. aureus, DAAT LL37 S. aureus or DAAT Indolicidin S. aureus at 128, 256 or 512 ng μL⁻¹ for 4 h, followed by serial dilution and plating on TSB agar. The PBS-treated control showed dense colony growth, indicating that S. aureus remained highly viable in the absence of DAAT treatment. In contrast, all three tail-fibre-retargeted DAAT formulations markedly reduced colony formation compared with the control group.

Representative plates showed that DAAT Cecropin, DAAT LL37 and DAAT Indolicidin treatment resulted in only scattered bacterial colonies, whereas the control plate was nearly covered with colonies. This difference indicated that tail-fibre-retargeted DAAT acquired direct killing activity against extracellular S. aureus. Among the three formulations, DAAT Cecropin and DAAT LL37 showed slightly fewer visible colonies than DAAT Indolicidin in the representative plates, but all DAAT-treated groups displayed a clear antibacterial effect. These results demonstrate that tail-fibre retargeting enabled DAAT particles to recognize and kill extracellular S. aureus, supporting their further use in intracellular infection and in vivo antibacterial treatment experiments.

### 3.6 DAAT promotes wound healing and tissue repair in a murine subcutaneous Staphylococcus aureus infection model

To evaluate the in vivo therapeutic effect of DAAT-mediated antimicrobial peptide delivery, a murine subcutaneous *Staphylococcus aureus* infection model was established. As shown in Fig. 4F, PBS-treated wounds exhibited persistent exudation, ulceration, and delayed healing, indicating sustained infection. In contrast, DAAT-Cecropin-*S. aureus*, DAAT-LL37-*S. aureus*, and DAAT-Indolicidin-*S. aureus* treatments markedly improved wound recovery, as shown by reduced exudate, progressive scab formation, and faster wound closure. By day 12, DAAT-treated wounds showed more complete surface repair than PBS-treated wounds.

Histological analysis further supported the wound-healing effect of DAAT treatment (Fig. 4G). H&E staining showed that DAAT-treated tissues had reduced inflammatory infiltration and improved epidermal structure compared with PBS-treated infected wounds. Masson’s trichrome staining revealed enhanced collagen deposition and more organized dermal remodeling in DAAT-treated groups. Together with reduced cytokine levels and decreased bacterial colonies, these results indicate that DAAT-mediated antimicrobial peptide delivery promotes infected wound healing by combining bacterial clearance, inflammatory alleviation, and tissue repair.

## 4. Discussion

Intracellular bacterial infection remains difficult to treat because pathogens can evade immune clearance, reduce exposure to antibacterial agents and persist as protected intracellular reservoirs after entering host cells (Peyrusson et al., 2020; Hommes et al., 2022; Volk et al., 2024). To address this delivery barrier, we developed Directed Antimicrobial Assault Technology (DAAT), a PVC nanosyringe-based antimicrobial peptide (AMP) delivery platform. By fusing AMPs with a Pdp1-derived loading module, peptide cargos were recruited into the PVC inner tube during particle assembly, integrating host-tolerant expression, protective encapsulation and intracellular injection delivery in one system (Jiang et al., 2022; Kreitz et al., 2023). This design converts AMPs, which are often difficult to express and deliver, into programmable nanosyringe payloads for intracellular bacterial therapy. The successful construction of DAAT-Cecropin, DAAT-LL37 and DAAT-Indolicidin demonstrates the modularity of this platform. TEM showed that all three particles retained PVC-like nanosyringe morphology, while SDS-PAGE indicated that AMP loading did not markedly disrupt the structural protein composition of PVC particles. HiBiT luminescence and low-molecular-weight cargo bands further confirmed successful AMP loading rather than nonspecific co-purification.

In the GFP-expressing USA300/Eno-GFP infection model, all three DAAT formulations suppressed intracellular bacterial growth in a dose-dependent manner. DAAT-LL37 showed the strongest activity, with an inhibition rate reaching 86.76%, whereas DAAT-Cecropin retained a broader effective range and DAAT-Indolicidin showed a narrower therapeutic window. Microscopy and colony spot-plating assays further supported reduced viable intracellular bacteria and alleviated infection-associated host-cell damage after DAAT treatment. This study also defined a practical working window for DAAT therapy. Although 512 ng μL⁻¹ showed strong antibacterial activity, it caused detectable stress in HT-29 cells (Xu et al., 2026). By contrast, 256 ng μL⁻¹ maintained antibacterial efficacy while better preserving host-cell morphology and viability, and was selected as the preferred working concentration. Repeated treatment at this dose increased live-cell numbers and promoted recovery of infected cells, indicating that DAAT reduces intracellular bacterial burden while preserving host-cell survival. Overall, this study establishes DAAT as a programmable PVC-derived AMP nanosyringe platform for intracellular bacterial therapy.

Tail-fibre modification gave DAAT an additional extracellular antibacterial function. By remodeling the tail-fibre targeting recognition domain toward *S. aureus*, DAAT could recognize Gram-positive bacteria, anchor to the bacterial surface and execute contractile injection-mediated killing (Xia et al., 2011; Li et al., 2016; Koç et al., 2016; Kreitz et al., 2023). The antibacterial spot-plating assay supported this design, showing that retargeted DAAT directly suppressed extracellular bacterial growth. This result expands DAAT beyond an intracellular AMP delivery carrier and suggests that PVC-derived particles can also function as active antibacterial nanomachines against extracellular bacteria. In this way, DAAT combines Pdp1-guided AMP loading for intracellular treatment with tail-fibre-mediated bacterial targeting for extracellular clearance, providing a coordinated strategy against the intracellular–extracellular escape cycle of *S. aureus* (Peyrusson et al., 2020; Hommes et al., 2022; Volk et al., 2024).

The therapeutic relevance of this dual-mode design was further supported in the mouse subcutaneous infection model, where intracellular AMP-delivering DAAT and extracellular *S. aureus*-targeting DAAT were administered together. DAAT treatment promoted more normal wound healing and markedly reduced the bacterial burden recovered from skin tissues, indicating effective antibacterial activity in a complex tissue environment. These in vivo results extend the value of DAAT beyond cell-culture assays and show that intracellular reservoir clearance and extracellular bacterial targeting can operate together as a synergistic therapeutic strategy. Together, the tail-fibre remodeling and mouse infection experiments establish DAAT as a programmable PVC-derived platform that integrates protected AMP delivery, direct extracellular bacterial attack and tissue-level infection control.

## 5. Conclusion and Outlook

This study establishes DAAT as a programmable PVC-derived antimicrobial peptide nanosyringe for synergistic intracellular and extracellular antibacterial therapy. Pdp1-guided loading enables host-tolerant AMP production, protective intraluminal encapsulation and targeted intracellular delivery, addressing key barriers of AMP expression, stability and access to host-protected bacteria. In parallel, remodeling the tail-fibre recognition domain redirects DAAT toward extracellular *S. aureus*, enabling bacterial binding, surface anchoring and contractile injection-mediated killing. Thus, DAAT combines intracellular reservoir clearance with extracellular bacterial targeting, expanding PVC-derived systems from protein delivery tools into adaptable precision antibacterial platforms.

The future value of DAAT lies in moving beyond natural antimicrobial peptides toward programmable, de novo-designed antibacterial cargos. Instead of relying only on existing AMP sequences, DAAT could be equipped with synthetic protein payloads engineered to attack intracellular bacteria through complementary mechanisms, such as direct bacterial damage, virulence suppression and immune-evasion disruption. This would transform DAAT from a peptide carrier into a modular antibacterial machine capable of executing multi-layered intervention inside protected infection reservoirs. At the same time, the thick cell wall of Gram-positive bacteria remains a physical barrier for contractile injection, making penetration efficiency a critical engineering frontier. Future optimization of DAAT recognition, anchoring and injection modules could ultimately create a new class of precision antibacterial nanomachines that do not simply deliver drugs, but actively seek, breach and dismantle persistent bacterial infections.

## RESOURCE AVAILABILITY

### Lead contact

Further information and requests for resources and reagents should be directed to and will be fulfilled by the lead contact, Xianchao Feng.

## Data and code availability

- All data supporting the findings of this study are available within the article and its supplementary information files.
- This paper does not report original code.
- Any additional information required for reanalyzing the data reported in this paper is available from the lead contact upon request.

## Author contributions

L. Feng. and Y. Qiao contributed to Investigation, Methodology, Data curation, Writing – original draft. H. Xu, G. Wang, S. Ren and X. Ouyang contributed to Software, Formal analysis and Conceptualization. N. Song, X. Zhao and X. Feng. contributed to Supervision, Resources, Funding acquisition, Writing – Review & Editing.

## Acknowledgments

This work was supported by the National Natural Science Foundation of China (grant nos. 32401252, 32172143 and 318732014), and the Key Research and Development Program of Shaanxi Province (Program No. 2024SF-ZDCYL-03-23). We are deeply grateful to all colleagues and collaborators for their valuable contributions, insightful discussions, and technical assistance throughout this study. We also sincerely acknowledge the broader community whose support facilitated the completion of this work. Finally, we respectfully honor the experimental animals whose sacrifices enabled the progress of this research, and we reaffirm our commitment to the highest standards of animal welfare and ethical research practices.

